# Transcriptional memory-like imprints and enhanced functional activity in γδ T cells following resolution of malaria infection

**DOI:** 10.1101/2020.05.05.078717

**Authors:** Rasika Kumarasingha, Lisa J. Ioannidis, Waruni Abeysekera, Stephanie Studniberg, Dinidu Wijesurendra, Ramin Mazhari, Daniel P. Poole, Ivo Mueller, Louis Schofield, Diana S. Hansen, Emily M. Eriksson

## Abstract

γδ T cells play an essential role in the immune response to malaria infection. However, long-lasting effects of malaria infection on the γδ T cell population still remain inadequately understood. This study investigated transcriptional changes and memory-like functional capacity of malaria pre-exposed γδ T cells using a *Plasmodium chabaudi* infection model. We show that multiple genes associated with effector function (chemokines, cytokines and cytotoxicity) and antigen-presentation were upregulated in *P. chabaudi*-exposed γδ T cells compared to γδ T cells from naïve mice. This transcriptional profile was positively correlated with profiles observed in conventional memory CD8^+^ T cells and was accompanied by enhanced reactivation upon secondary encounter with *Plasmodium*-infected red blood cells *in vitro*. Collectively our data demonstrate that *Plasmodium* exposure result in “memory-like imprints” in the γδ T cell population and also promotes γδ T cells that can support antigen-presentation during subsequent infections.

## Introduction

γδ T cells are unconventional T cells that display characteristic features of both innate and adaptive immunity. Their capacity to respond rapidly to non-peptide antigens in an MHC-independent manner places them as part of the innate first line of defense against numerous pathogens. Additionally, emerging evidence supports the concept that γδ T cells also display memory T cell-like abilities. This includes prolonged recall responses upon reinfection in various disease and vaccine models, which contribute to protective immunity ^1–6^. Recent studies have now started to delineate a more in-depth understanding of these adaptive-like γδ T cells. For example, it has been described that the TCR of tissue-resident γδ T cells has an intrinsic ability to distinguish between distinct antigen-stimulus and in this way promote either clonal or non-clonal responses ^7^ whereas adaptive-like γδ T cells found in peripheral human blood are suggested to be restricted to specific subsets of the γδ T cell population ^8^.

*Plasmodium* infection, which is responsible for the induction of malaria in humans, elicits a multifaceted response activating a wide range of immune cells, including γδ T cells. Extensive evidence shows that γδ T cells are part of the immediate innate response during human malaria infection where they are found to be cytotoxically active and produce cytokines associated with both protective immunity and symptomatic episodes ^9–15^. The underlying mechanisms by which γδ T cells either contribute to beneficial outcomes in the host or mediate pathogenesis remain to be fully elucidated. However, studies using field samples indicate that repeated exposure to the parasite promotes γδ T cells that are associated with the development of clinical symptoms ^13–15^ in an infected individual. In contrast, γδ T cells have been proposed to be part of the IFNγ-producing memory pool found upon re-exposure in experimentally controlled-malaria infected individuals ^16^.

In addition to human infections, γδ T cells are also highly involved in the immune response to murine malaria. In mice, they are a major source of cytokines and contribute to parasite clearance ^17–22^ and are essential for protective immunity following vaccination ^23^. This makes murine malaria infection models a useful platform to explore fundamental immunological questions related to *Plasmodium* infection and also enables the investigation of immune populations within various tissues not readily available from infected individuals. *P. chabaudi* infection in C57BL/6 mice is a self-resolving infection, and this infection model has been used to successfully elucidate various aspects of γδ T cell biology. γδ T cells proliferate extensively in response to *P. chabaudi* infection and mice lacking γδ T cells experience exacerbated parasitemia ^21, 24–26^. More recently, γδ T cells from chronically infected mice were described to produce inflammatory chemokines such as CCL3 and CCL5 and also importantly m-CSF, which was vital to the control of recrudescence ^19^ suggesting that “antigen-experienced” γδ T cells play a role in the suppression of parasitemia in chronic infection. Although these studies further emphasize that γδ T cells are readily activated during acute *Plasmodium* infection, the lasting effect that *Plasmodium* exposure has on these cells and how this shapes the γδ T cell population is still inadequately understood. Consequently, we used the *P. chabaudi* murine malaria infection model to investigate transcriptional profiles of γδ T cells from naïve and malaria-exposed mice, 12 weeks after completion of anti-malarial drug treatment. Our findings revealed that antigen-experienced γδ T cells display a transcriptional profile that shares features with that of conventional memory CD8^+^ T cells and have enhanced functional capacity. Thus, our data support the notion that γδ T cells differentiate and acquire a memory-like phenotype after infection. These observations advance our basic understanding of unconventional T cell biology in malaria but may also give insight into how responses to vaccines or other pathogens can be affected by malaria pre-exposure.

## Results

### Increased frequencies of multifunctional γδ T cells in drug-cured *P. chabaudi*-exposed mice

The hallmark of memory T cells is increased functional capacity upon secondary encounter with specific antigen, which commonly includes IFNγ production and cytotoxic activity. To establish whether similar responses were generated in γδ T cells following *Plasmodium* infection, we compared responses of naïve and pre-exposed γδ T cells upon antigen re-encounter. Since spleen and liver are central to the immune response to *P. chabaudi* infection ^27, 28^ and are organs that have previously been shown to contain tissue resident innate memory cells ^29, 30^, we assessed γδ T cell responses in both of these organs. To that end, C57BL/6 mice were infected with *P. chabaudi* and drug-cured on day 14 post-infection (p.i.) to clear parasitemia completely. Twelve weeks after completion of drug-treatment spleens and livers were harvested (Figure 1A). Splenocytes and liver lymphocytes were subsequently isolated and stimulated *in vitro* with *P. chabaudi-*infected red blood cells (iRBC) or uninfected RBC (uRBC) as background controls. Cells from naïve mice were included to measure baseline responses. After a 24 h incubation, CD107a surface expression (as a measure of cytotoxic activity) and IFNγ production were assessed by flow cytometry (Figure 1B). We found that a significantly higher frequency of γδ T cells that were both CD107a^+^ and produced IFNγ were present in the spleens of previously infected mice compared to naïve mice (Figure 1C, P< 0.0001). No significant differences were observed with γδ T cells that produced only IFNγ or were CD107a^+^. Similarly, no significant differences in functionality were detected in the liver-derived γδ T cells from pre-exposed *P. chabaudi*-infected mice and naïve mice (Figure 1D). This showed that *P. chabaudi* infection resulted in the induction of multi-functional memory-like γδ T cells.

**Figure 1.**
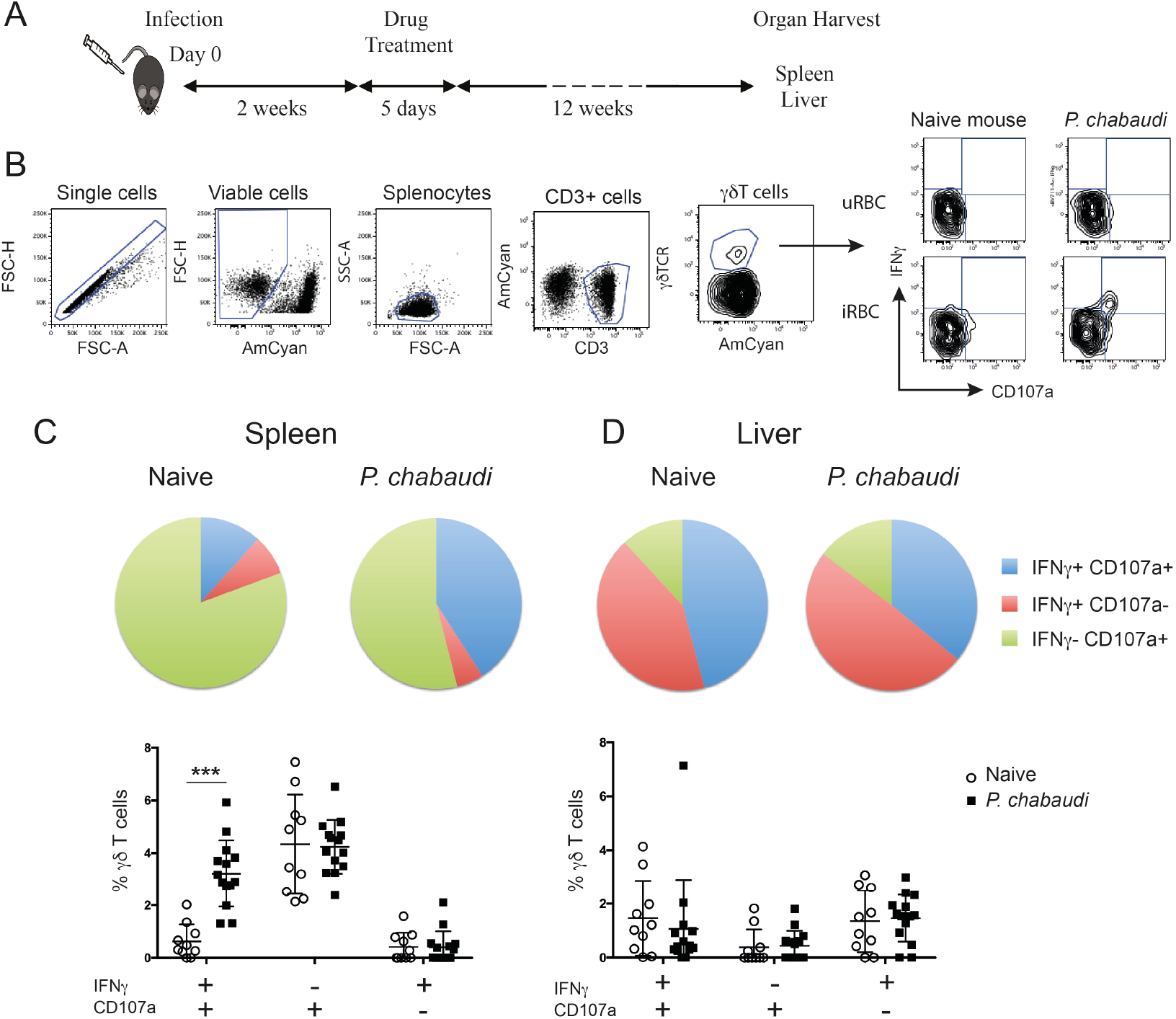
Increased frequency of IFNγ^+^CD107a^+^ γδ T cells in previously infected mice. A) C57BL/6 mice were infected with *P. chabaudi* and then drug-treated with chloroquine and pyrimethamine two weeks later. Twelve weeks following completion of drug-treatment cells were isolated and stimulated with iRBCs or uRBCs and frequencies of IFNγ^+^ and/or CD107a^+^ cells were assessed. B) Representative flow cytometry plots illustrating the gating strategy. Frequencies of IFNγ^+^ and/or CD107a^+^ C) splenocytes and D) liver lymphocytes from previously infected mice (*P. chabaudi* black squares, n=14) and naïve control (white circles, n=10) after stimulation. In the pie chart the data are presented as the frequency of IFNγ^+^ CD107a^+^ (blue), IFNγ^+^ CD107a^−^ (red) and IFNγ^−^ CD107a^+^ (green) γδ T cells in each group following uRBC background subtraction. The data in the scatter plot are presented as mean ± SD following uRBC background subtraction. The data represent two independent experiments. Statistical analysis was performed using Student’s t-tests. ***P < 0.001

### Responding γδ T cells express an effector memory-like phenotype

Previous studies indicate that the γδ T cells that provide effector functions during acute malaria infection express surface markers that resemble conventional αβ T effector memory cells ^16, 19^. To assess the phenotype of the responding γδ T cells of previously exposed mice after full resolution of infection, we stimulated spleen-derived γδ T cells from drug-treated mice or naïve mice *in vitro* and stained the cells for the surface markers CD62L and CD44. This enabled the γδ T cells to be subdivided into CD62L^+^CD44^−^ naïve cells, CD62L^+^CD44^+^ central memory cells (CM) and CD62L^−^CD44^+^ effector memory cells (EM; Figure 2A). The frequency of IFNγ^+^CD107a^+^ double positive γδ T cells in each subset was assessed in both groups of mice. Representative flow cytometry plots of these responses are presented in Figure 2B. Upon stimulation with iRBC, responding γδ T cells were found to predominantly express an EM phenotype and frequencies of IFNγ^+^CD107a^+^ EM γδ T cells were significantly higher in previously *P. chabaudi*-infected mice compared to naïve control mice (P< 0.0001; Figure 2C). This demonstrated that γδ T memory-like responses were specifically confined within the EM subset. Furthermore, the increase in frequency of responding cells did not reflect an overall increase of EM γδ T cells in the pre-exposed mice as assessment of the γδ T cell composition showed no differences in frequencies (Figure 2D) or cell numbers (Figure 2E) of naïve, CM or EM γδ T cells between *P. chabaudi* exposed mice and uninfected controls.

**Figure 2.**
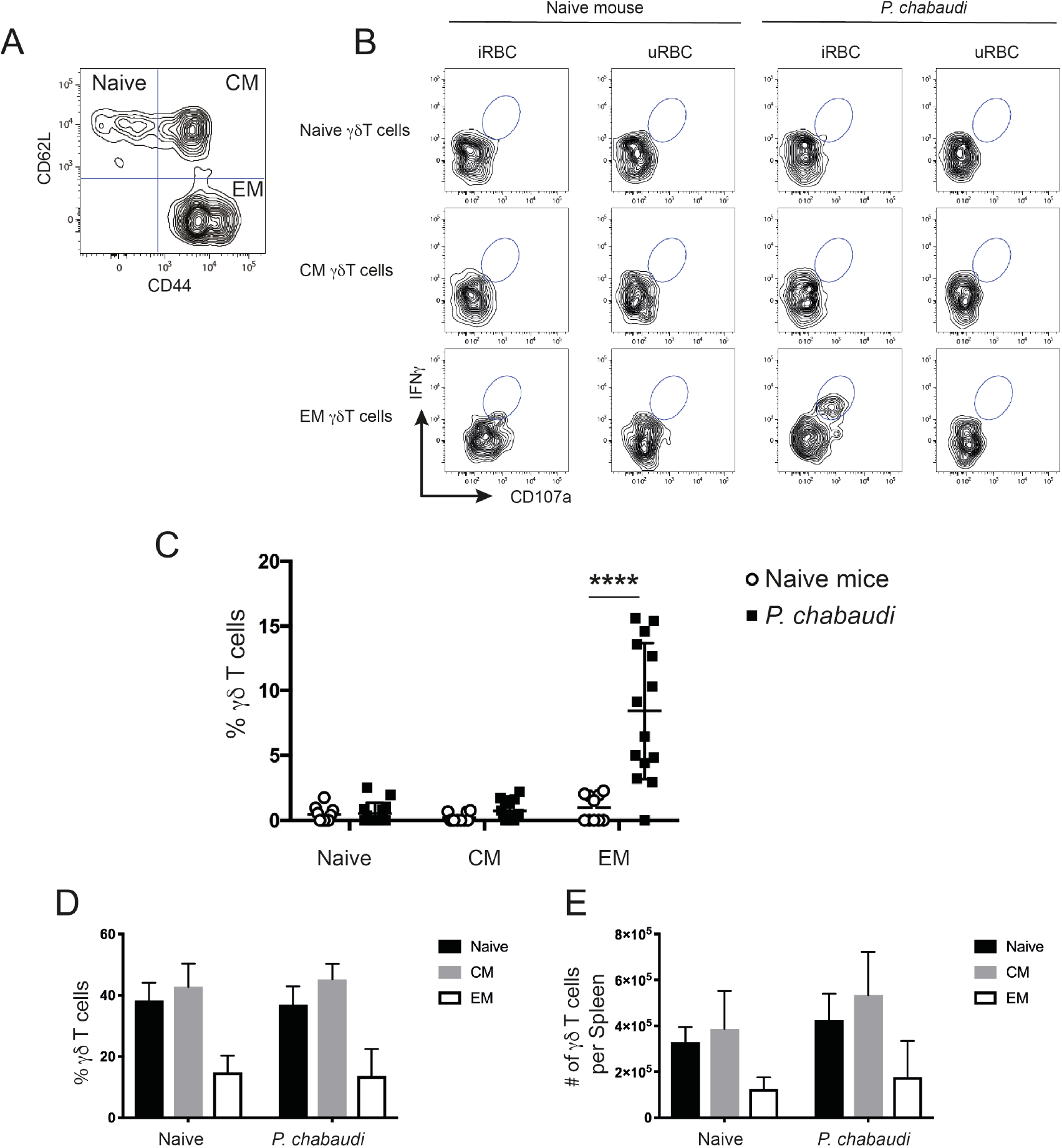
*In vitro* re-stimulated and activated γδ T cells express CD44, but lack CD62L expression. Splenocytes from previously infected and drug treated mice and naïve controls were restimulated *in vitro* with iRBC or uRBCs. Representative contour plot to A) distinguish between CD62L^+^CD44^−^ (Naïve), CD62L^+^CD44^+^ (CM) and CD62L^−^CD44^+^ EM γδ T cells. B) Representative contour plots showing frequency of IFNγ^+^CD107a^+^ γδ T cells for each subset after 24 h stimulation with either iRBC or uRBC from naïve or *P. chabaudi* pre-exposed mice. C) Summary of IFNγ^+^CD107a^+^ naïve, CM and EM γδ T cells after iRBC stimulation following subtraction of background levels determined from uRBC stimulations in previously *P. chabaudi*-infected mice (filled squares; n=14) and naïve controls (open circles; n=10). Overall D) frequency and E) number of γδ T cells per spleen of naïve, CM and EM γδ T cells (mean±SD) in naïve or *P. chabaudi* pre-exposed mice. The data represent two independent experiments. Statistical analysis was performed using C-E) Student’s t-tests ****P < 0.0001

### Transcriptional profile changes in EM γδ T cells from drug-treated *P. chabaudi* exposed mice compared to EM γδ T cells from naïve mice

We have shown that γδ T cells expressing an EM-phenotype are re-activated upon re-encounter with *P. chabaudi* iRBC in previously infected mice (Figure 2). As the frequency and number of EM γδ T cells in the spleens were not different between the naïve control group and the pre-exposed mice, this indicated that this memory-like enhanced responsiveness was due to intrinsic changes of the cells. To investigate this, EM γδ T cells were FACS-sorted from mice 12 weeks after they had been drug treated to clear *P. chabaudi* infection (n=5) and from naïve mice (n=5; Figure 3) and RNA-sequencing was used to examine transcriptional profiles. A total of 207 differentially expressed (DE) genes in *P. chabaudi* pre-exposed EM γδ T cells compared to EM γδ T cells from naïve mice were observed relative to a fold change threshold of 1.5 (Supplemental Table 1). Expression levels and log-fold changes were plotted in a Mean-Difference (MD) plot (Figure 3A) of which 96 genes were significantly upregulated (indicated in red) and 111 genes were significantly down regulated (indicated in blue). The upregulated genes included MHC class II-related genes (H2-Dmb2 and H2-A) and also IFNγ and NKg7, which corresponded to the observed functional phenotype of enhanced IFNγ production and cytotoxicity in the pre-exposed EM γδ T cells (Figure 2). The chemokine genes (CCL3, CCL4 and CCL5) were also upregulated in these memory-like γδ T cells, which is similar to what had previously been reported to be upregulated in γδ T cells during an active infection (Mamedov 2018). Cytokine receptor genes (Il1r and Il23r), scavenger-receptor gene (Cd163l1) and transcription factor gene (Sox13) were among the down regulated genes. The top 75 DE genes are summarized in a heatmap presenting up- and down regulated genes in each mouse (Figure 3B). Collectively, this shows that malaria-infection causes significant transcriptional changes in the EM γδ T cell population, which is still observed in absence of an active infection.

**Figure 3.**
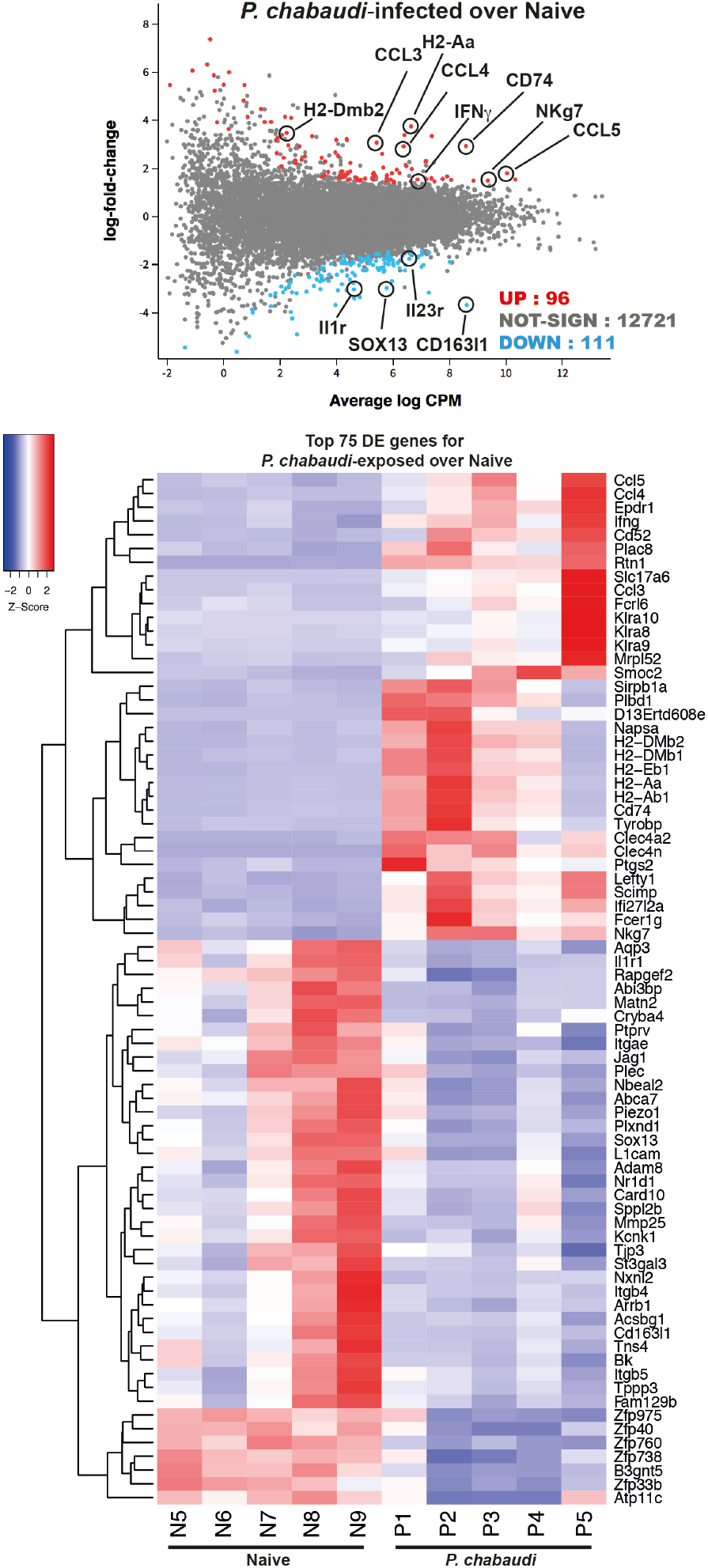
RNA-sequencing of EM γδ T cells from *P. chabaudi* pre-exposed mice and naïve controls. EM γδ T cells from drug-treated naïve mice (n=5 donors) and *P. chabaudi* pre-exposed mice (n=5) were FACS sorted followed by RNA extraction and RNA-sequencing. Differential gene expression for *P. chabaudi* over naïve mice was summarized in A) mean-difference (MD) plot of log2 expression fold-changes against the average log-expressions for each gene. The differentially expressed (DE) genes relative to a fold change threshold of 1.5 are highlighted, with points colored in red and blue indicating up- and down regulated genes respectively. B) Heatmap of the expressions of the top 75 DE genes between *P. chabaudi* and naïve mice. Each vertical column represents genes for each mouse. For a given gene the red and blue coloring indicates increased and decreased expression in *P. chabaudi* compared to naïve respectively.

### Genes involved in antigen presentation and processing are upregulated in pre-exposed EM γδ T cells

To understand the biological processes affected by previous exposure to malaria in the EM γδ T cell population, gene ontology (GO) pathway analysis was performed. Among the twenty most highly enriched GO terms in the upregulated biological processes, 7 were associated with antigen-processing and presentation. In addition, genes were enriched for processes involving positive regulation of acute inflammatory responses and response to IFNγ (Supplemental Figure 1A). Barcode plots and bar plots illustrating the enrichment of all genes in selected pathways showed that antigen-processing and presentation pathway included upregulation of MHC class II-related genes (H2-Aa, H2-Dmb2, H2-Ab1, H2-Eb1, H2-Dmb1), genes that support antigen-processing and presentation (Clec4a2, Flt3, Cd74, Ifng) and genes for FC receptor expression (Fcrgr2b, Fcer1g; Figure 4A). There were also enrichment of genes that suggested an increased responsiveness to IFNγ stimulation as shown by upregulation of chemokine and cytokine genes (Ccl3, Ccl4, Ccl5, Ifng, Xcl1), interferon induced transmembrane protein genes (Ifitm2, Ifitm3) and MHC class II-related genes (H2-Aa, H2-Ab1, H2-Eb1), but down regulation of IL23r (Figure 4B). In addition, gene enrichment analysis suggested that pre-exposed EM γδ T cells have the potential to contribute to a sustained inflammatory response as shown by upregulation of Fcer1a, Alox5bp. Ptgs2, Fcer1g and Ccl5 combined with down regulation of Adam8 (Figure 4C).

**Figure 4.**
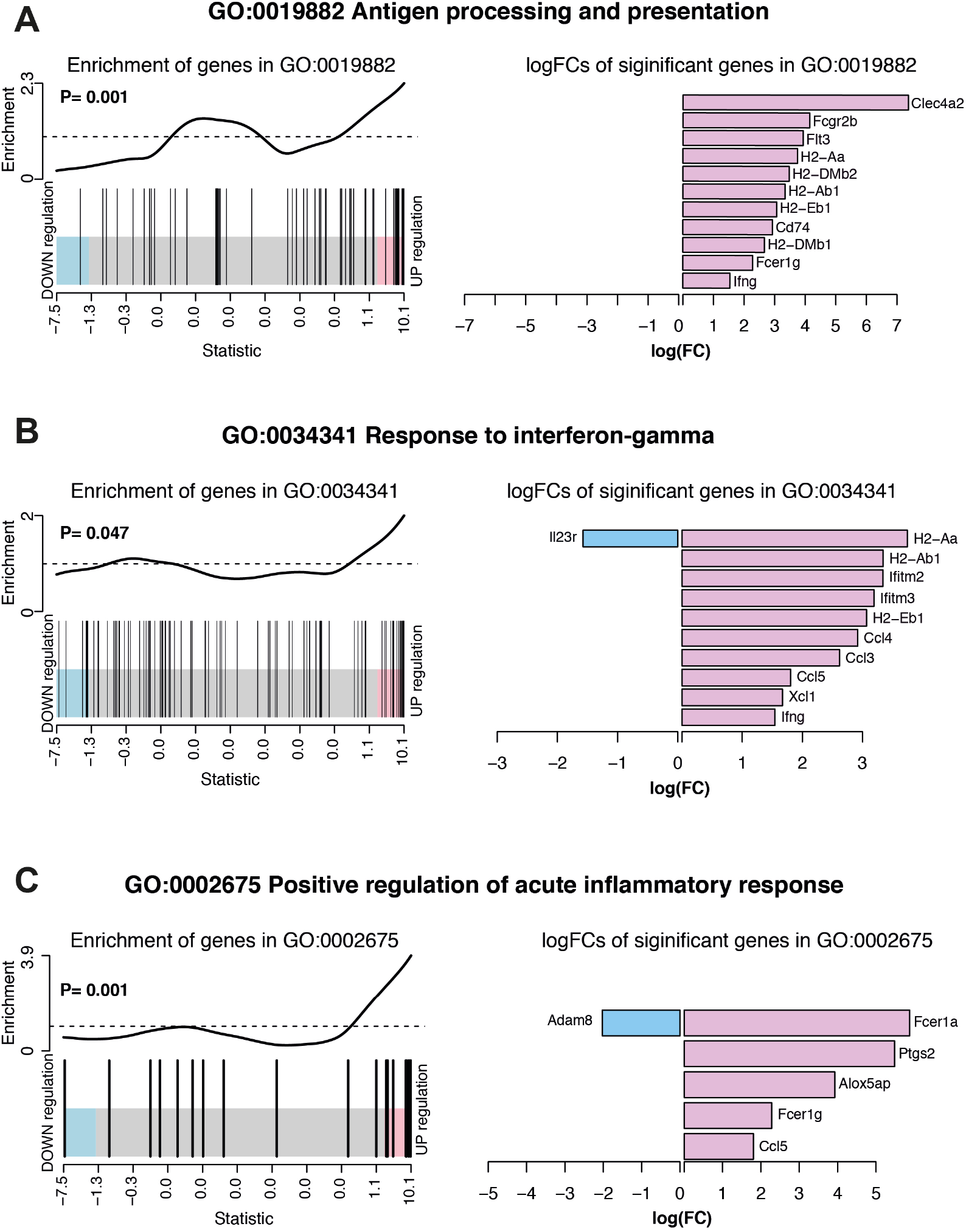
Summary of significant up or down regulated genes in selected upregulated biological processes. Barcode plots for enrichment of the pathway genes along with p-values relative to gene enrichment tested using ROAST method (left panel) and bar graphs of log fold changes of the significant pathway genes (right panel) for pathways A) GO:0019882 antigen processing and presentation, B) GO: 0034341 response to IFNγ and C) GO:0002675 positive regulation of acute inflammatory response. The barcode plot ranks genes right to left from most up-to most down regulated in *P. chabaudi* mice, with genes in the pathways marked by vertical bars. The bar graph show log fold changes of significantly upregulated and down regulated genes in the pathway using pink bars and blue bars respectively.

Some of the most significantly down regulated biological processes included cell-substrate adhesion and cellular response to stimulus (Supplemental Figure 1B). Considering that responsiveness to IFNγ stimulation was increased (Supplemental Figure 1A), decrease in the biological process of cellular response to stimulus suggests that the pre-exposed EM γδ T cell population is modulated to only respond to specific conditions such as presence of IFNγ. Barcode plots and bar plots illustrating the enrichment of all genes in these down regulated pathways showed that a total of 85 DE genes were represented in the cellular response to stimulus (Figure 5A). The 3 most down regulated genes in this pathway were Itgb4, Plxnd1 and Tspan2, which are all associated with signal transduction and cell-cell signaling. The most upregulated gene in this pathway was Fcer1a, which has been associated with an immune suppressive role in APCs ^31^. The enrichment of all genes in the cell-substrate adhesion pathway included down regulated integrin genes (Itgb4, Itga5, Itgb5), protein kinases (Trmp7, Slk) and genes associated with cell recruitment, adhesion and migration (Adam8, Jag1, Lamc1, L1cam) whereas Epdr and Smoc2 genes were upregulated (Figure 5B).

**Figure 5.**
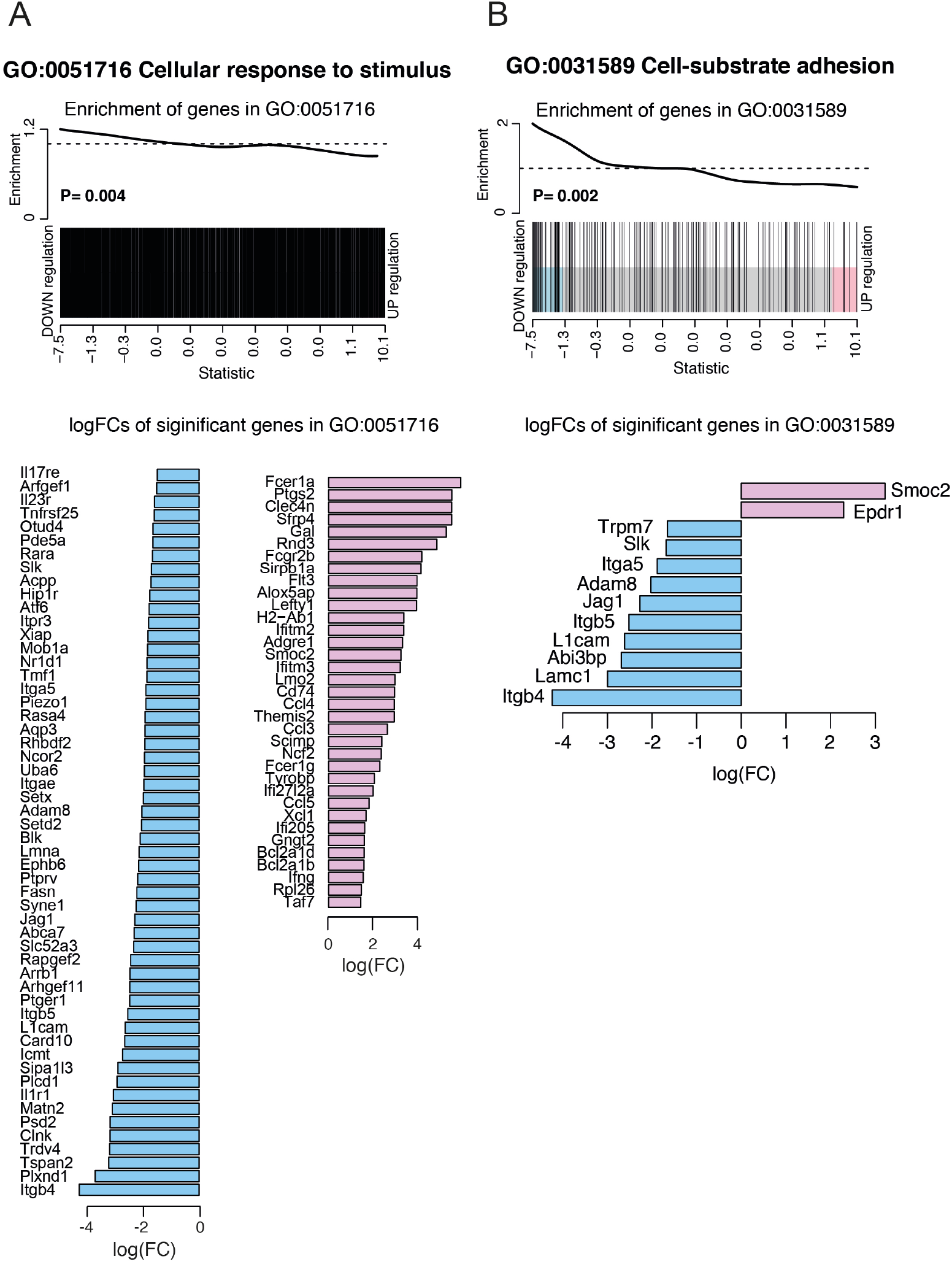
Summary of significant up or down regulated genes in selected down regulated biological processes. Barcode plots for enrichment of the pathway genes along with p values relative to gene enrichment tested using ROAST method (top panel) and bar graphs of log fold changes of the significant pathway genes (bottom panel) for pathways A) GO:0051716 cellular response to stimulus and B) GO: 0031589 cell-substrate adhesion. The barcode plot ranks genes right to left from most up-to most down regulated in *P. chabaudi* mice, with genes in the pathways marked by vertical bars. The bar graph show log fold changes of significantly upregulated and down regulated genes in the pathway using pink bars and blue bars respectively.

### Differentially expressed genes in pre-exposed EM γδ T cells are positively correlated with differentially expressed genes in resting memory CD8^+^ T cells

A previous study demonstrated that conventional CD8^+^ memory T cells have distinct transcriptional profiles that significantly differ from those of their naïve counterparts ^32^. To examine whether the transcriptional profile observed in the pre-exposed γδ T cells resembled that of conventional memory T cells, we compared our DE expression data (Supplemental Table 1) with a previously described signature defining resting CD8^+^ memory T cells ^32^ (Russ et al. Supplemental Table 2). A total of 43 DE genes were represented in both gene sets, of which 32 were upregulated and 11 were down regulated DE genes (Figure 6A). These overlapping genes presented in a heatmap (Figure 6B) included genes that were associated with hallmark functions of conventional memory T cells such as cytokine/chemokine production and cytotoxicity (Ccl4, Ccl5, Ccl3, Ifng, Nkg7). Genes involved in antigen presentation and processing (Clec4a2, Fcgr2b, H2-Aa, H2-Dmb2, H2-Ab1, H2-Eb1, Cd74, H2-Dmb1, Fcer1g), which was a prominent transcriptional signature of the memory-like EM γδ T cell DE gene set, also overlapped with the DE genes from CD8^+^ memory T cells. Furthermore, enrichment analysis showed that both up and down regulated DE genes in the EM γδ T cell gene set positively correlated with the DE genes in the CD8^+^ memory T cell gene set (P= 0.008; Figure 6C). Altogether, this supports the novel concept that *Plasmodium* exposure induces EM γδ T cells with a transcriptional profile resembling conventional memory T cells.

**Figure 6.**
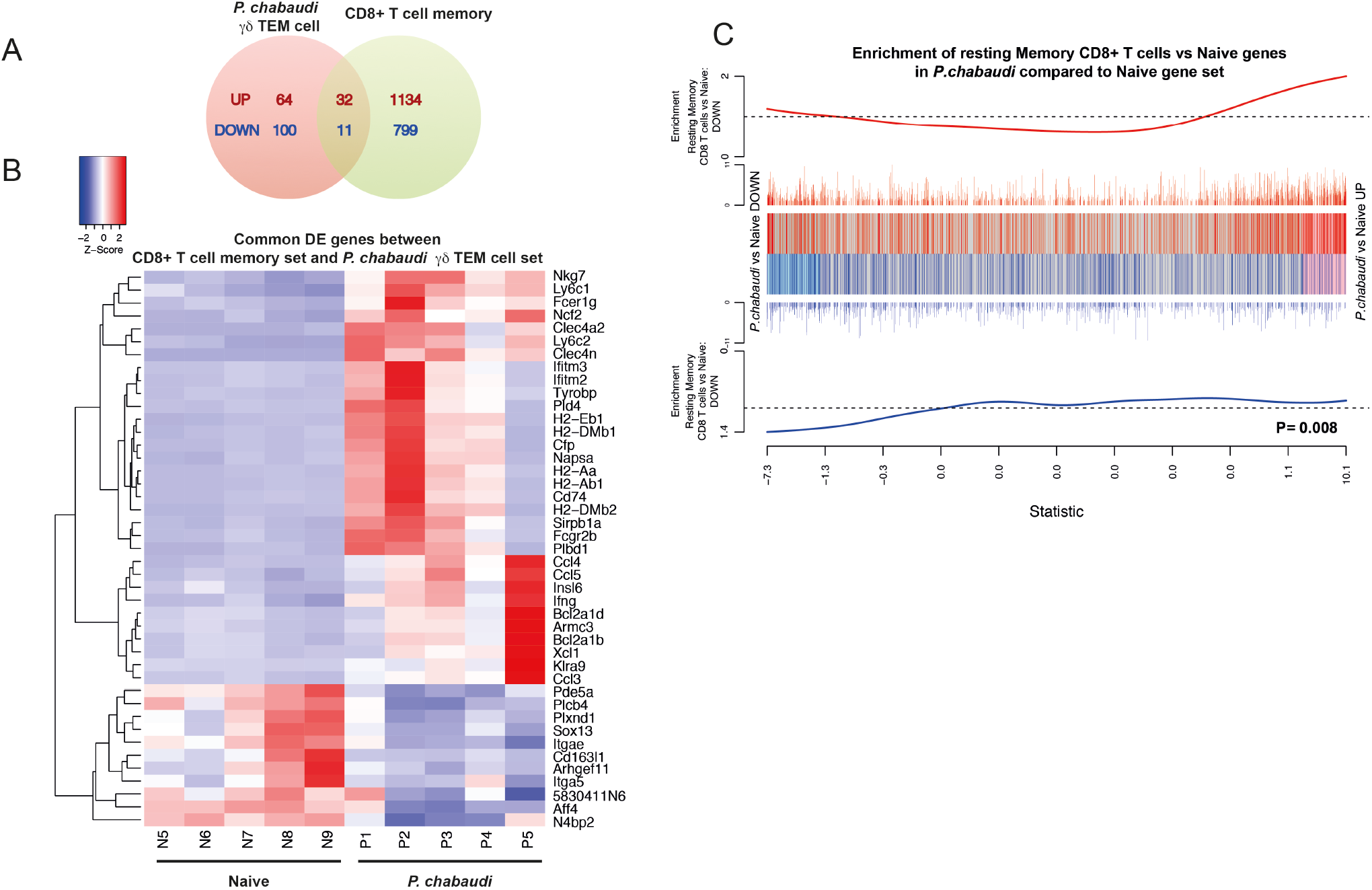
Differentially expressed genes in *P. chabaudi* pre-exposed γδ T cells are positively correlated with differentially expressed genes in CD8^+^ memory T cells. A) Venn diagram showing the number of overlapping and non-overlapping up-regulated (red) and down regulated (blue) genes. B) Heatmap of the gene expression relative to *P. chabaudi* pre-exposed γδ T cells data for the genes commonly significantly regulated (overlapping DE genes) between *P. chabaudi* pre-exposed γδ T cells data and CD8^+^ memory T cell data. Each vertical column represents genes for each mouse. For a given gene the red and blue coloring indicates increased and decreased expression in *P. chabaudi* compared to naïve respectively. C) Barcodeplot for the enrichment of DE genes in the resting CD8^+^ memory T cell data in *P. chabaudi* compared to naïve in the *P. chabaudi* pre-exposed γδ T cell data, along with the ROAST p-value for the gene set testing.

## Discussion

Immunological memory is considered a definitive feature of adaptive immunity, allowing clonal selection of antigen-specific T and B lymphocytes where differentiation to memory cells is the basis for enhanced secondary responses to subsequent challenge. In contrast, innate immunity was long considered to lack such memory features and instead rely on a repertoire of invariant receptors. However, recent studies demonstrate that a number of cells of the innate immune system have the capacity to establish immunological memory. Emerging functional evidence suggest that this includes γδ T cells in specific disease models ^2, 33^. However, whether exposure to antigen induces transcriptional profiles in EM γδ T cells associated with enhanced effector function have remained elusive. In this study we used a malaria infection model to understand whether “memory-like imprints” were detectable in γδ T cells after the infection was cleared and whether this was associated with memory-like γδ T cell responses. We found that the transcriptional profile in pre-exposed EM γδ T cells was significantly different from EM γδ T cells from naïve mice and that differentially expressed genes in the pre-exposed EM γδ T cells were positively correlated with previously reported differentially expressed genes in resting CD8^+^ memory T cells. This showed that although γδ T cell populations in both naïve mice and previously *P. chabaudi*-infected mice were classified as “memory” populations based on traditional surface markers, only pre-exposed γδ T cells were observed to resemble that of conventional T cell memory.

Consistent with their memory-like transcriptional profile, we also found that pre-exposure to antigen resulted in enhanced functional capacity of responding γδ T cells upon encounter with cognate antigen. It has been suggested that as γδ T cells emerge from the thymus, they have already acquired a functional imprint, which limits their plasticity in the periphery ^34–36^. Furthermore, functionally distinct γδ T cells seem to have specific tissue distribution where spleen-derived γδ T cells are predominately prone to producing IFNγ ^34^. We found that pre-exposed γδ T cells were multifunctional as they produced both IFNγ and were cytotoxically active. However, the EM γδ T cell population previously-exposed to malaria displayed significant reductions in the expression of genes associated with IL-17 responses, suggesting limitation to their functional plasticity after *Plasmodium* infection. Apart from low gene expression of IL-17a, this included significantly lower expression levels of *Sox13* and *Il1r1* genes. Sox13 is a lineage specific γδ T cell transcription factor ^37^, which promotes IL-17 producing γδ T cells ^38^ and IL-1 has recently been indicated to play an important role in supporting IL-17 production by antigen-specific T cells *in vivo.* Cells from *Il1r1*-deficient mice had dramatically reduced IL-17 production compared to cells from wild-type mice ^39^. Furthermore IL-17 producing γδ T cells have been shown to rapidly respond to IL-23, which induces and supports IL-17 production ^40–42^. Interestingly following *Plasmodium* exposure, EM γδ T cells have down regulated their *Il-23r* gene expression suggesting that they are less responsive to endogenous IL-23. This indicates that the functionally intrinsic characteristics of EM γδ T cells is altered with infection and is maintained in absence of parasites.

We showed here that the γδ T cell population in the spleen not only acquires memory-like characteristics, but also potentially fill an additional role as APCs. Although antigen-presentation and processing by γδ T cells has previous been described, this characteristic remains relatively unexplored. This function is seemingly acquired upon TCR activation and human Vδ2 T cells activated with isopentenyl pyrophosphate induced high levels of APC-related molecules, which resulted in a functional capacity to present antigens to αβ T cells ^43^. In *P. falciparum*-infected individuals there is an increase of Vγ9Vδ2 T cells that express APC-related surface markers and this expression was induced by iRBCs ^44^. These cells were also able to elicit αβ T cell responses *in vitro* suggesting that γδ T cells may simply supplement existing APC populations. However, spleen-derived γδ T cells reside in an organ that plays a central role in the capacity to control and clear parasites and are in a location that allows them to encounter and remove blood-borne antigens and also initiate innate and adaptive immune responses. It is possible that following an initial malaria infection once an adaptive memory has been established, exposed γδ T cells promote specific adaptive T cell functions. In support of this proposition, intestinal γδ T cells have been found to have APC function and elicit distinct CD4^+^ T cell responses compared to responses induced by typical professional APCs ^45^. A comprehensive understanding of the APC-capacities of tissue-resident γδ T cells and the specific functions that they provide for subsequent *Plasmodium* infections remains to be determined.

The work presented here demonstrates that blood-stage *Plasmodium* infection has a profound effect on the splenic γδ T cell population, modifying its response capacity and gene expression profile. While our observations here support the existence of traditional memory cells with augmented secondary responses upon antigen re-encounter, it also appears that this functional capacity may not necessarily be the only role for these cells. Our findings here suggest a model by which antigen-experienced γδ T cells undergo transcriptional changes that allows them to fulfil a novel role as antigen-presenting cells in subsequent infections. These findings have important implications for our understanding of the role of γδ T cells in host immunity and gives insight into potential therapeutic modulations that can be achieved.

## Material and Methods

### Mice and Mouse infection

Female C57BL/6 mice aged 6-8 weeks were infected with 5×10^4^ *Plasmodium chabaudi* iRBC intravenously. All mice were drug-treated on day 14 p.i. with an intraperitoneal injection of chloroquine (CQ; 10mg/kg) and pyrimethamine (10mg/kg) followed by CQ (0.6 mg/ml) and pyrimethamine (70 μg/ml) containing water for 5 days. Spleens and livers were removed 12 weeks after completion of drug treatment. The experimental design is summarized in Figure 1A. Organs from drug-treated naïve mice were used as controls. All procedures involving mice were approved by the Walter and Eliza Hall Institute animal ethics committee (2015.020).

### *In vitro* cell stimulation

Single cell suspensions from spleen or liver were prepared as previously described ^46^. Wholeblood from *P. chabaudi-*infected donors were obtained during the dark cycle to obtain mature parasites ^47^. The blood was washed in RPMI and 0.5-1 ml of blood in medium was overlayed onto 12.17 ml of a 74% percoll gradient as described in ^48^ and centrifuged at 5000 g for 20 min at room temperature. IRBCs were collected from the interface and washed with culture medium. Isolated iRBCs were co-incubated with splenocytes and liver lymphocytes at a ratio of 1:1 for 24 h. Brefeldin A (Sigma, St. Louis, MO) and GolgiStop (BD Biosciences, San Jose, CA) were added for the final 8 h of incubation.

### Flow Cytometry and FACS sorting

A total of 1×10^6^ splenocytes or liver lymphocytes were surface stained with Brilliant Violet (BV) 421-conjugated anti-CD107a (clone 1D4B, BioLegend, San Diego, CA) during the 24 h stimulation. Further surface staining following stimulation was performed with antibody mixtures in FACS buffer (phosphate buffer saline containing 0.5% bovine serum albumin (BSA) and 2mM ethylenediaminetetraacetic acid (EDTA) on ice for 30 min. Antibodies used included: Fluorescein isothiocyanate (FITC)-conjugated anti-CD3 (clone 145-2C11), PerCP Cy5.5-conjugated anti-γδTCR (clone GL3), allophycocyanin (APC)-conjugated anti-CD27 (clone LG.3A10), (all from BioLegend), Alexa700-conjugated anti-CD44 (clone IM7) and Brilliant Violet (BV) 605-conjugated anti-CD62L (clone MEL-14, BD Biosciences, San Jose, CA). Aqua live/dead amine reactive dye (Life Technologies, Carlsbad, CA) was used for dead cell exclusion. Intracellular staining was performed after 2% paraformaldehyde fixation and permeabilization with Perm 2 buffer (BD Biosciences) using BV711-conjugated anti-IFNγ (clone XMG1.2, BioLegend). Samples were analyzed on a customized four-laser Fortessa flow cytometer (BD Biosciences). Data analysis was performed using FlowJo 9.9.6 software (TreeStar, Ashland, OR) and Boolean gating. For FACS sorting, splenocytes were surface stained with CD3, γδTCR, CD62L and CD44 as above to identify and collect γδ T cells with a phenotype associated with T effector memory (EM, CD62L^−^ CD44^+^).

### Library preparation and transcriptome sequencing

EM γδ T cells from 5 naïve control mice and 5 mice that had been previously infected with *P. chabaudi* and then drug-treated to clear the infections were FACS sorted. Total RNA was isolated from sorted cells using the Isolate II RNA mini kit (Bioline, London, UK) according to manufacturer’s instructions. RNA was quantified using the Agilent TapeStation 2200 system (Santa Clara, CA). An input of 1 ng of total RNA were prepared and indexed separately for sequencing using the CloneTech SMART ultra-low RNA input Prep Kit (Illumina, San Diego, CA) as per manufacturer’s instruction. The indexed libraries were pooled and diluted to 1.5pM for paired end sequencing (2× 76 cycles) on a NextSeq 500 instrument using the v2 150 cycle High Output kit (Illumina) as per manufacturer’s instructions. The base calling and quality scoring were determined using Real-Time Analysis on board software v2.4.6, while the FASTQ file generation and de-multiplexing utilized bcl2fastq conversion software v2.15.0.4. Paired-end 75bp. Between 16 and 56 million read pairs were generated for each sample and reads were aligned to the *Mus musculus* genome (mm10) using the Subread aligner ^49^. The number of read pairs overlapping each mouse Entrez gene was summarized using featureCount ^50^ and Subread’s built-in NCBI gene annotation. Genes were filtered using filterByExpr function in edgeR ^51^ software package. Genes without current annotation and Immunoglobulin genes were also filtered. Differential expression (DE) analysis was undertaken using the edgeR and limma ^52^ software packages. Library sizes were normalized using the trimmed mean of M-values (TMM) method ^53^. Log2 fold-changes were computed using voom ^54^. Differential expression was assessed relative to a fold change threshold of 1.5 using the TREAT ^55^ function, a robust empirical Bayes procedure ^56^ implemented in the limma package. The false discovery rate (FDR) was controlled below 0.05 using the method of Benjamini and Hochberg ^57^. Over-representation of Gene Ontology (GO) terms for the differentially expressed genes was identified using the goana function in limma package. Barcode plots illustrating the enrichment of interested pathway genes were drawn using the barcode plot function in limma package ^58^.

### Statistical Analysis

Statistical analyses were performed using Prism 8.0 (GraphPad software, San Diego, CA) Flow cytometry data was analyzed using the Student’s t-test. Statistical significance was considered P ≤ 0.05

## Acknowledgements

We wish to thank Liana Mackiewicz and Carolina Alvarado at WEHI for technical assistance.

## Author contribution

RK and LJI performed experiments and critically reviewed the manuscript, WA and DPP analysed data and critically reviewed the manuscript, DW, RM and SS analysed data, IM, DSH and LS provided conceptual input into the study design and critically reviewed the manuscript, EME conceived and performed experiments, analysed data and prepared the manuscript.

