## Supplemental Figure 1 for "Transcriptional memory-like imprints and enhanced functional activity in γδ T cells following resolution of malaria infection"

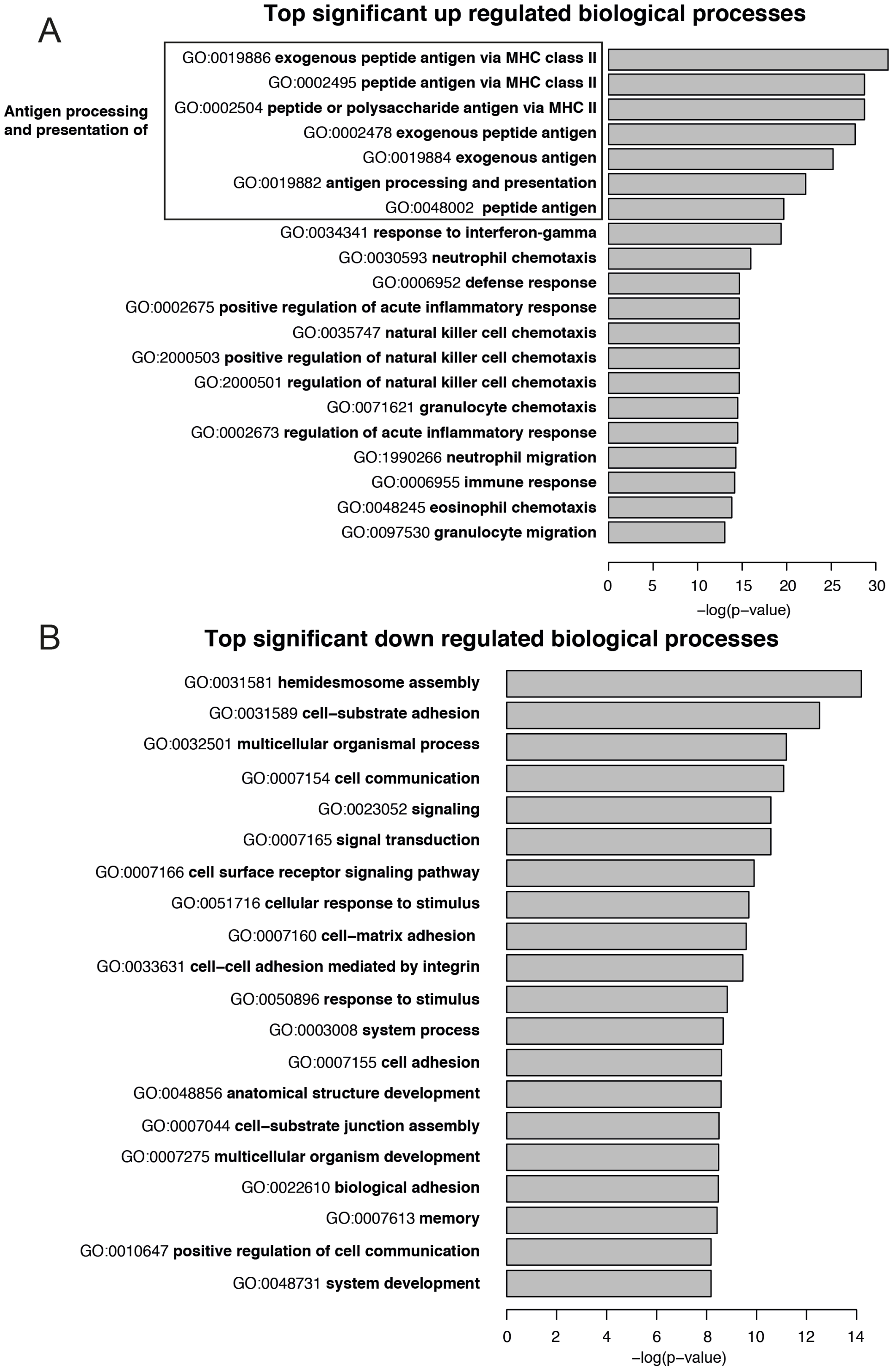


**Supplemental Figure 1. Pathway analysis of differentially expressed genes.** Gene Ontology (GO) terms for the differentially expressed genes were identified and the top 20 significantly A) upregulated and B) down regulated biological processes are presented.
