## Supplemental Table 1 for "Transcriptional memory-like imprints and enhanced functional activity in γδ T cells following resolution of malaria infection"

### Supplemental Table 1. Differentially expressed genes in resting EM γδ T cells for *P. chabaudi* exposed mice over naïve mice relative to a fold change threshold of 1.5.

| GeneID | Symbol | logFC | P.Value | adj.P.Val |
|---|---|---|---|---|
| 16640 | Klra9 | 3.39044649 | 1.42E-08 | 0.00018394 |
| 20302 | Ccl3 | 2.61819118 | 3.59E-08 | 0.00023202 |
| 140919 | Slc17a6 | 8.58584287 | 1.29E-07 | 0.00055684 |
| 20303 | Ccl4 | 2.92039006 | 2.31E-07 | 0.00074802 |
| 17181 | Matn2 | -3.0656979 | 7.88E-07 | 0.00175484 |
| 20668 | Sox13 | -2.9775554 | 8.14E-07 | 0.00175484 |
| 14961 | H2-Ab1 | 3.34151569 | 1.29E-06 | 0.00209603 |
| 14969 | H2-Eb1 | 3.06629838 | 1.47E-06 | 0.00209603 |
| 16639 | Klra8 | 3.39858743 | 1.58E-06 | 0.00209603 |
| 244233 | Cd163l1 | -3.6734629 | 1.63E-06 | 0.00209603 |
| 192897 | Itgb4 | -4.2356539 | 1.78E-06 | 0.00209603 |
| 104001 | Rtn1 | 6.32010768 | 2.01E-06 | 0.00210703 |
| 320712 | Abi3bp | -2.687706 | 2.24E-06 | 0.00210703 |
| 14960 | H2-Aa | 3.74438646 | 2.28E-06 | 0.00210703 |
| 67971 | Tppp3 | -3.3576541 | 3.87E-06 | 0.00333899 |
| 67784 | Plxnd1 | -3.6726208 | 5.56E-06 | 0.00445659 |
| 76933 | Ifi27l2a | 1.9771099 | 5.86E-06 | 0.00445659 |
| 16149 | Cd74 | 2.93113707 | 7.00E-06 | 0.00484936 |
| 16407 | Itgae | -1.9508531 | 7.15E-06 | 0.00484936 |
| 16525 | Kcnk1 | -2.2320014 | 7.98E-06 | 0.00484936 |
| 108105 | B3gnt5 | -2.1318702 | 8.00E-06 | 0.00484936 |
| 75124 | Nxnl2 | -3.9675443 | 8.25E-06 | 0.00484936 |
| 64074 | Smoc2 | 3.22005352 | 9.93E-06 | 0.00558654 |
| 320940 | Atp11c | -3.0596644 | 1.18E-05 | 0.0063155 |
| 14127 | Fcer1g | 2.28114455 | 1.22E-05 | 0.0063155 |
| 22700 | Zfp40 | -2.8183597 | 1.34E-05 | 0.00667617 |
| 16628 | Klra10 | 3.17360354 | 1.52E-05 | 0.00728868 |
| 56620 | Clec4n | 5.47291964 | 1.61E-05 | 0.00737075 |
| 16728 | L1cam | -2.6150969 | 1.70E-05 | 0.00737075 |
| 16177 | Il1r1 | -3.0255573 | 1.71E-05 | 0.00737075 |
| 94180 | Acsbg1 | -2.1520114 | 1.78E-05 | 0.00742548 |
| 434179 | Zfp975 | -2.7566189 | 1.93E-05 | 0.00781934 |
| 12143 | Blk | -2.0813922 | 2.03E-05 | 0.00786837 |
| 109689 | Arrb1 | -2.4470744 | 2.12E-05 | 0.00786837 |
| 105298 | Epdr1 | 2.29149016 | 2.13E-05 | 0.00786837 |
| 66857 | Plbd1 | 3.1880755 | 2.23E-05 | 0.00800138 |
| 27403 | Abca7 | -2.2961181 | 2.44E-05 | 0.00853689 |
| 19225 | Ptgs2 | 5.48449503 | 2.92E-05 | 0.00951065 |
| 240034 | Zfp760 | -2.6778093 | 2.94E-05 | 0.00951065 |
| 20304 | Ccl5 | 1.80542457 | 2.94E-05 | 0.00951065 |
| 12959 | Cryba4 | -3.1216169 | 3.33E-05 | 0.0104986 |
| 217169 | Tns4 | -3.4001073 | 3.66E-05 | 0.0111969 |
| 320832 | Sirpb1a | 4.11075448 | 3.72E-05 | 0.0111969 |
| 15000 | H2-DMb2 | 3.4752987 | 4.52E-05 | 0.01299018 |
| 217166 | Nr1d1 | -1.8385267 | 4.55E-05 | 0.01299018 |
| 23833 | Cd52 | 1.54680848 | 4.62E-05 | 0.01299018 |
| 408068 | Zfp738 | -2.231938 | 5.04E-05 | 0.01382819 |
| 20441 | St3gal3 | -2.4250712 | 5.16E-05 | 0.01382819 |
| 16541 | Napsa | 1.82045567 | 5.24E-05 | 0.01382819 |
| 73218 | Sppl2b | -1.9845924 | 5.47E-05 | 0.01415938 |
| 22177 | Tyrobp | 2.03778719 | 5.85E-05 | 0.01470045 |
| 69623 | Zfp33b | -2.2383166 | 6.00E-05 | 0.01470045 |
| 327957 | Scimp | 2.36750583 | 6.24E-05 | 0.01470045 |
| 231507 | Plac8 | 2.10617985 | 6.26E-05 | 0.01470045 |
| 27375 | Tjp3 | -1.9661036 | 6.27E-05 | 0.01470045 |
| 18810 | Plec | -2.0152577 | 6.45E-05 | 0.01470045 |
| 13924 | Ptprv | -2.1715927 | 6.55E-05 | 0.01470045 |
| 52673 | D13Ertd608e | 6.07348703 | 6.59E-05 | 0.01470045 |
| 105844 | Card10 | -2.6295919 | 6.85E-05 | 0.0147293 |
| 16449 | Jag1 | -2.2705059 | 6.91E-05 | 0.0147293 |
| 26888 | Clec4a2 | 7.36981245 | 6.95E-05 | 0.0147293 |
| 11828 | Aqp3 | -1.9149296 | 7.31E-05 | 0.0150935 |
| 72310 | Nkg7 | 1.49537211 | 7.35E-05 | 0.0150935 |
| 227737 | Fam129b | -2.4930972 | 7.68E-05 | 0.01551876 |
| 16419 | Itgb5 | -2.518283 | 8.26E-05 | 0.01635881 |
| 240047 | Mmp25 | -2.021048 | 8.35E-05 | 0.01635881 |
| 68836 | Mrpl52 | 1.71245827 | 8.63E-05 | 0.0166606 |
| 677296 | Fcrl6 | 2.52556764 | 9.08E-05 | 0.01714184 |
| 76089 | Rapgef2 | -2.4179422 | 9.26E-05 | 0.01714184 |
| 14999 | H2-DMb1 | 2.6663498 | 9.30E-05 | 0.01714184 |
| 235627 | Nbeal2 | -2.2397252 | 9.41E-05 | 0.01714184 |
| 13590 | Lefty1 | 3.91201824 | 9.83E-05 | 0.01766426 |
| 15978 | Ifng | 1.54412634 | 0.00010009 | 0.01773421 |
| 234839 | Piezo1 | -1.8827456 | 0.00010328 | 0.01788065 |
| 11501 | Adam8 | -2.024776 | 0.00010561 | 0.01788065 |
| 19216 | Ptger1 | -2.4559706 | 0.00010627 | 0.01788065 |
| 17067 | Ly6c1 | 1.77670531 | 0.00010645 | 0.01788065 |
| 74691 | Tdrd9 | -4.894266 | 0.00011056 | 0.01806239 |
| 64009 | Syne1 | -2.2244327 | 0.00011247 | 0.01806239 |
| 57294 | Rps27 | 1.81403551 | 0.00011273 | 0.01806239 |
| 545389 | Cep170 | -2.0176272 | 0.00011312 | 0.01806239 |
| 235626 | Setd2 | -2.0350638 | 0.00011845 | 0.01868366 |
| 13733 | Adgre1 | 3.28728561 | 0.00012078 | 0.01881755 |
| 69698 | Slc52a3 | -2.3118547 | 0.00012221 | 0.01881755 |
| 69781 | 1600010M07 | 6.00329433 | 0.00013267 | 0.02018718 |
| 100041546 | Ly6c2 | 1.52389971 | 0.00013728 | 0.02064608 |
| 224697 | Adamts10 | -1.7600819 | 0.00014343 | 0.02130377 |
| 269254 | Setx | -1.9703498 | 0.00014495 | 0.02130377 |
| 74194 | Rnd3 | 4.81909751 | 0.00014927 | 0.02169261 |
| 12045 | Bcl2a1b | 1.56732395 | 0.00015629 | 0.0220893 |
| 16905 | Lmna | -2.1187115 | 0.00015995 | 0.0220893 |
| 217082 | Hlf | -2.6080009 | 0.00016201 | 0.0220893 |
| 12047 | Bcl2a1d | 1.58363573 | 0.00016267 | 0.0220893 |
| 546643 | I830127L07R | 2.64517731 | 0.00016386 | 0.0220893 |
| 231380 | Uba6 | -1.9325607 | 0.00016425 | 0.0220893 |
| 20379 | Sfrp4 | 5.47280312 | 0.00016729 | 0.0220893 |
| 213498 | Arhgef11 | -2.4534256 | 0.0001683 | 0.0220893 |
| 19401 | Rara | -1.6487025 | 0.00016845 | 0.0220893 |
| 11958 | Atp5k | 1.71041274 | 0.00017328 | 0.0220893 |
| 547023 | Gm6014 | 1.59037474 | 0.00017408 | 0.0220893 |
| 100038467 | A630072L19I | -3.1879796 | 0.00017412 | 0.0220893 |
| 104759 | Pld4 | 2.23339953 | 0.0001742 | 0.0220893 |
| 20335 | Sec61g | 1.71316333 | 0.00018335 | 0.02285935 |
| 66141 | Ifitm3 | 3.1891748 | 0.00018393 | 0.02285935 |
| 16963 | Xcl1 | 1.67046898 | 0.00018558 | 0.02285935 |
| 333789 | N4bp2 | -2.1499791 | 0.00020016 | 0.0244229 |
| 93736 | Aff4 | -1.6704461 | 0.00020374 | 0.02462745 |
| 18636 | Cfp | 2.5068555 | 0.00021903 | 0.0261136 |
| 19724 | Rfx1 | -1.7988245 | 0.00022007 | 0.0261136 |
| 99031 | Osbpl6 | -5.4399794 | 0.00022644 | 0.02650221 |
| 242474 | Tmem245 | -1.9398074 | 0.00022926 | 0.02650221 |
| 105203 | Fam208b | -2.1364652 | 0.00022949 | 0.02650221 |
| 14419 | Gal | 5.23497664 | 0.0002367 | 0.02703276 |
| 27278 | Clnk | -3.1480376 | 0.00023855 | 0.02703276 |
| 106064 | AW549877 | -1.4509389 | 0.00024242 | 0.02703276 |
| 58800 | Trpm7 | -1.6551586 | 0.00024546 | 0.02703276 |
| 100114901 | Trdv4 | -3.1632699 | 0.0002466 | 0.02703276 |
| 546336 | Prrg1 | -3.7274869 | 0.00024663 | 0.02703276 |
| 100042480 | Nhsl2 | -2.3384695 | 0.00024908 | 0.02707175 |
| 70882 | Armc3 | 2.09645514 | 0.00025411 | 0.02738847 |
| 212281 | Zfp729a | -1.8037739 | 0.00026404 | 0.02822362 |
| 11305 | Abca2 | -2.1059443 | 0.00026962 | 0.02849805 |
| 433771 | Minos1 | 1.46099902 | 0.00027444 | 0.02849805 |
| 18798 | Plcb4 | -1.8991147 | 0.00027573 | 0.02849805 |
| 668548 | Gm9234 | 1.5886041 | 0.00027762 | 0.02849805 |
| 19942 | Rpl27 | 1.49528932 | 0.000279 | 0.02849805 |
| 13848 | Ephb6 | -2.1407699 | 0.0002806 | 0.02849805 |
| 623286 | Gm6415 | 1.73980233 | 0.00028276 | 0.02849805 |
| 64380 | Ms4a4c | 1.56184576 | 0.0002846 | 0.02849805 |
| 18799 | Plcd1 | -2.9029124 | 0.00028877 | 0.02849805 |
| 232157 | Mob1a | -1.8235198 | 0.00029138 | 0.02849805 |
| 100310809 | Gm10509 | -1.9455024 | 0.00029467 | 0.02849805 |
| 85030 | Tnfrsf25 | -1.5834613 | 0.00029471 | 0.02849805 |
| 20874 | Slk | -1.6840805 | 0.00029525 | 0.02849805 |
| 80876 | Ifitm2 | 3.33832857 | 0.00030793 | 0.02936592 |
| 70747 | Tspan2 | -3.1933786 | 0.00031025 | 0.02936592 |
| 217344 | Rhbdf2 | -1.9191638 | 0.00031105 | 0.02936592 |
| 17970 | Ncf2 | 2.34277338 | 0.00032008 | 0.02985542 |
| 74206 | Sipa1l3 | -2.870379 | 0.00032085 | 0.02985542 |
| 57295 | Icmt | -2.7002589 | 0.00032518 | 0.03004208 |
| 103012 | Firre | -2.885473 | 0.00033104 | 0.03036672 |
| 235682 | Zfp445 | -1.6368334 | 0.00033843 | 0.03065132 |
| 664903 | Rps15a-ps4 | 1.52584779 | 0.00033889 | 0.03065132 |
| 71679 | Atp5h | 1.46354865 | 0.00034347 | 0.03085009 |
| 595139 | E030024N20 | 1.9564233 | 0.00035041 | 0.03105678 |
| 54127 | Rps28 | 1.63489106 | 0.00035057 | 0.03105678 |
| 442834 | D830031N03 | -5.6264778 | 0.00035918 | 0.03160264 |
| 232286 | Tmf1 | -1.841913 | 0.00037438 | 0.03271751 |
| 209590 | Il23r | -1.5742096 | 0.00038148 | 0.03311473 |
| 14255 | Flt3 | 3.93307654 | 0.00039181 | 0.03343625 |
| 68052 | Rps13 | 1.52797482 | 0.00039619 | 0.03343625 |
| 76486 | Ly6k | 2.43981225 | 0.00039855 | 0.03343625 |
| 56318 | Acpp | -1.7027226 | 0.00039856 | 0.03343625 |
| 233115 | Dpy19l3 | -1.8132615 | 0.00039868 | 0.03343625 |
| 244234 | 5830411N06 | -1.8919018 | 0.00040359 | 0.03343625 |
| 14710 | Gngt2 | 1.59211673 | 0.00040515 | 0.03343625 |
| 226691 | Ifi207 | 4.14276731 | 0.00040587 | 0.03343625 |
| 14934 | Gypa | 4.47460481 | 0.00042532 | 0.03481664 |
| 27762 | Vwa7 | -3.8726915 | 0.00044147 | 0.03564666 |
| 100217422 | Snord13 | 2.4390167 | 0.00044415 | 0.03564666 |
| 70081 | Zfp995 | -2.0314871 | 0.0004466 | 0.03564666 |
| 11798 | Xiap | -1.8115982 | 0.0004494 | 0.03564666 |
| 240753 | Plekha6 | -3.2662489 | 0.00044954 | 0.03564666 |
| 66475 | Rps23 | 1.53622924 | 0.00045199 | 0.03564666 |
| 218850 | Fam208a | -1.7662213 | 0.00047048 | 0.03688001 |
| 100043695 | Gm15483 | 1.65788997 | 0.00048099 | 0.03714473 |
| 29816 | Hip1r | -1.7443726 | 0.00048107 | 0.03714473 |
| 268373 | Ppia | 1.49147232 | 0.00048247 | 0.03714473 |
| 27356 | Insl6 | 1.77847073 | 0.00049245 | 0.03768816 |
| 226695 | Ifi205 | 1.60727328 | 0.00050021 | 0.03784678 |
| 24074 | Taf7 | 1.42555474 | 0.00050037 | 0.03784678 |
| 268449 | Rpl23a | 1.57644911 | 0.00051487 | 0.03871673 |
| 72672 | Zfp518a | -1.5869731 | 0.00051826 | 0.03874669 |
| 14125 | Fcer1a | 5.87666897 | 0.00054134 | 0.04023936 |
| 100043882 | Gm14277 | 1.55978207 | 0.00054489 | 0.0402722 |
| 67671 | Rpl38 | 1.87136115 | 0.00054916 | 0.04035677 |
| 211673 | Arfgef1 | -1.4996966 | 0.00055826 | 0.04079382 |
| 16440 | Itpr3 | -1.7847781 | 0.0005751 | 0.04178855 |
| 100043296 | Gm4342 | 1.47870869 | 0.00057988 | 0.0419007 |
| 245671 | Klf8 | -4.9847596 | 0.00058537 | 0.04206218 |
| 54153 | Rasa4 | -1.9113718 | 0.00060161 | 0.04283607 |
| 77700 | 9130208D14 | -2.4595416 | 0.00060277 | 0.04283607 |
| 73656 | Ms4a6c | 1.80779298 | 0.00060755 | 0.04294023 |
| 240038 | Zfp994 | -1.7955281 | 0.00061668 | 0.04334875 |
| 381605 | Tbc1d2 | -3.9868935 | 0.00062098 | 0.04341456 |
| 100042335 | Rps15a-ps5 | 1.50810763 | 0.00062536 | 0.04348578 |
| 230787 | Themis2 | 2.91912847 | 0.00063685 | 0.04378543 |
| 14130 | Fcgr2b | 4.14352871 | 0.00064223 | 0.04378543 |
| 633188 | Gm20762 | 1.53664557 | 0.00064238 | 0.04378543 |
| 100124677 | Trbv13-3 | 1.88412132 | 0.00064321 | 0.04378543 |
| 226519 | Lamc1 | -2.9979216 | 0.0006607 | 0.04451856 |
| 73945 | Otud4 | -1.6303022 | 0.00066086 | 0.04451856 |
| 100042173 | Rps15a-ps6 | 1.64503684 | 0.00066518 | 0.04457726 |
| 14104 | Fasn | -2.1972034 | 0.0006725 | 0.04483534 |
| 667090 | Gm8451 | 1.6598696 | 0.0006883 | 0.04547373 |
| 16402 | Itga5 | -1.8821888 | 0.00068999 | 0.04547373 |
| 20602 | Ncor2 | -1.9287641 | 0.00069262 | 0.04547373 |
| 242202 | Pde5a | -1.6322369 | 0.00069731 | 0.04555069 |
| 11690 | Alox5ap | 3.92836672 | 0.00070132 | 0.04558242 |
| 213121 | Ankrd35 | -2.7322028 | 0.00072014 | 0.0465715 |
| 319565 | Syne2 | -1.6112692 | 0.00073176 | 0.04683652 |
| 100416706 | Zfp729b | -1.5515733 | 0.00073552 | 0.04683652 |
| 19861 | Rnu3b4 | 1.65181887 | 0.00073606 | 0.04683652 |
| 19941 | Rpl26 | 1.45725058 | 0.00073872 | 0.04683652 |
| 268670 | Zfp759 | -3.0108621 | 0.00078001 | 0.04914534 |
| 11941 | Atp2b2 | 3.64003229 | 0.0007833 | 0.04914534 |
| 16909 | Lmo2 | 2.95502143 | 0.00078654 | 0.04914534 |
| 57890 | Il17re | -1.4806967 | 0.00079281 | 0.04929907 |
| 74002 | Psd2 | -3.1439883 | 0.00079855 | 0.04941853 |
| 100134990 | Selenok-ps1 | 1.71535575 | 0.00080446 | 0.04954681 |
| 19850 | Rnu3a | 2.22061571 | 0.00081795 | 0.0499763 |
| 226641 | Atf6 | -1.767882 | 0.00082072 | 0.0499763 |
| 100039988 | Gm11826 | 1.85151395 | 0.00082302 | 0.0499763 |
